# PAI-1 Deficiency Drives Pulmonary Vascular Smooth Muscle Remodeling and Pulmonary Hypertension

**DOI:** 10.1101/2023.09.21.558893

**Authors:** Tatiana V. Kudryashova, Sergei Zaitsev, Lifeng Jiang, Benjamin J Buckley, Joshua P. McGuckin, Dmitry Goncharov, Iryna Zhyvylo, Derek Lin, Geoffrey Newcomb, Bryce Piper, Srimathi Bogamuwa, Aisha Saiyed, Leyla Teos, Marie Ranson, Paul J. Wolters, Michael J. Kelso, Mortimer Poncz, Horace M. DeLisser, Douglas B. Cines, Elena A. Goncharova, Laszlo Farkas, Victoria Stepanova

## Abstract

Pulmonary arterial hypertension (PAH) is a progressive and potentially a rapidly fatal disease characterized by vasoconstriction and remodeling of small pulmonary arteries (PA) leading to increased pulmonary vascular resistance and right heart failure. Central to the remodeling process is a switch of the smooth muscle cells in small PAs (PASMC) to a proliferative, apoptosis-resistant phenotype.

There is reason to suspect that the plasminogen activator system may play an important role in the remodeling program in PAH based on its roles in vascular post-injury restenosis, fibrosis, angiogenesis and tumorigenesis. Plasminogen activator inhibitor-1 (PAI-1) is the primary physiological inhibitor of the plasminogen activators - urokinase-type and tissue-type (uPA and tPA, respectively). Immunohistochemical and immunoblot analyses revealed that PAI-1 was deficient in smooth muscle areas of small remodeled PAs and early-passage PASMC from subjects with PAH compared to non-PAH controls. *PAI1-/-* male and female mice developed spontaneous pulmonary vascular remodeling and pulmonary hypertension (PH) as evidenced by significant increase in PA medial thickness, systolic right ventricular pressure, and right ventricular hypertrophy. Lastly, the uPA inhibitors upamostat (WX-671) and amiloride analog BB2-30F down-regulated mTORC1 and SMAD3, restored PAI-1 levels, reduced proliferation, and induced apoptosis in human PAH PASMC. We examined the effect of inhibition of uPA catalytic activity by BB2-30F on the development of SU5416/Hypoxia (SuHx)-induced PH in mice. Vehicle-treated SuHx-exposed mice had up-regulated mTORC1 in small PAs, developed pulmonary vascular remodeling and PH, as evidenced by significant increase of PA MT, sRVP, RV hypertrophy, and a significant decrease in the pulmonary artery acceleration time/pulmonary ejection time (PAAT/PET) ratio compared to age- and sex-matched normoxia controls, whereas BB2-30F-treated group was protected from all these pathological changes. Taken together, our data strongly suggest that PAI-1 down- regulation in PASMC from human PAH lungs promotes PASMC hyper-proliferation, remodeling, and spontaneous PH due to unopposed uPA activation. Further studies are needed to determine the potential benefits of targeting the PAI-1/uPA imbalance to attenuate the progression and/or reverse pulmonary vascular remodeling and PH.

## RESEARCH LETTER

The sentinel pathologic findings in pulmonary arterial hypertension (PAH) involve a pulmonary vasculopathy characterized by vasoconstriction and pathologic remodeling of the distal pulmonary arteries (PAs), which lead to luminal narrowing, increased PA pressure, right ventricular afterload, and potentially premature death due to right heart failure(1). Available treatment approaches, predominantly targeting the vasoconstriction component by modulating endothelin, nitric oxide and prostacyclin signaling pathways, do not reverse pulmonary vascular remodeling and/or fully reduce disease progression. There is a clear need for newer therapies based on molecular mechanisms focused on vascular remodeling to complement contemporary management(1).

Unchecked hyper-proliferation and resistance to apoptosis of distal PA smooth muscle cells (PASMC) are important pathological components of pulmonary vascular remodeling. However, the molecular mechanisms underlying these events are incompletely understood. Multiple factors, including genetic predisposition, epigenetic mechanisms, oxidative stress, metabolic reprograming, extracellular matrix remodeling, inflammation, and immune cell infiltration contribute to PASMC hyper-proliferation, remodeling, and PAH(2). Mechanistically, dysregulation of the bone morphogenic protein receptor 2 (BMPR2)/transforming growth factor-β (TGF-β) network and over-activation of pro-proliferative receptor tyrosine kinase and growth factors-mediated signaling pathways, including Akt/mechanistic target of rapamycin (mTOR), appear to play important roles in supporting the pathogenic PASMC phenotype and pulmonary vascular remodeling in PAH(1). The mechanistic link between these key molecular players in PAH is not fully established, and the strategies to reverse PASMC molecular re-programing, remodeling and PAH are under development(1, 2).

There is reason to suspect that the plasminogen activator system may play an important role in the remodeling program in PAH based on its roles in vascular post-injury restenosis, fibrosis, angiogenesis and tumorigenesis. Plasminogen activator inhibitor-1 (PAI-1) is the primary physiological inhibitor of the plasminogen activators - urokinase-type and tissue-type (uPA and tPA, respectively). PAI-1 interacts with Arg-346 within the enzymatically active site of two-chain uPA (tcuPA), inactivating the protease and forming a covalent uPA-PAI-1 complex(3), which is internalized by the low density lipoprotein (LDL) receptor related protein-1 (LRP-1) and degraded in lysosomes(4). PAI-1 deficiency leads to unopposed uPA activity, increased plasminogen activation and plasmin-mediated proteolysis(5), and activation of TGF-β. To date, both pro-proliferative(6) and anti-proliferative(7) roles for PAI-1 in PASMCs have been proposed, so the potential attractiveness of targeting PAI-1 signaling in PAH is not clear.

Here, we report that PAI-1 is deficient in PASMCs from small muscular PAs in PAH lungs. Furthermore, we show that loss of PAI-1 results in uncontested activation of uPA, spontaneous remodeling of pulmonary vasculature, and pulmonary hypertension (PH). Lastly, we report that pharmacological inhibition of uPA restores PAI-1, down-regulates both TGF-β/Smad3 and Akt/mTOR, suppresses proliferation and induces apoptosis in human PAH PASMCs, and halts pulmonary vascular remodeling and experimental PH in mice.

Using immunohistochemical and immunoblot analyses, we detected a marked decrease in PAI-1 protein *in situ* in smooth muscle α-actin (SMA)-positive areas in small, remodeled PAs from patients with PAH (**Fig. 1A**). PAI-1 was also decreased *in vitro* in distal early-passage human PAH PASMC compared to non-diseased controls **(Fig. 1B)**, accompanied by a significant increase in plasminogen activation **(Fig. 1C)** and cell proliferation **(Fig. 1D)**. siRNA-induced silencing of PAI-1 in PASMCs from non-diseased (control) subjects led to a significant increase of mTORC1-specific phosphorylation of ribosomal protein S6 (**Fig. 1E**), suggesting the link between PAI-1 deficiency and PASMC proliferation.

**Fig. 1.**
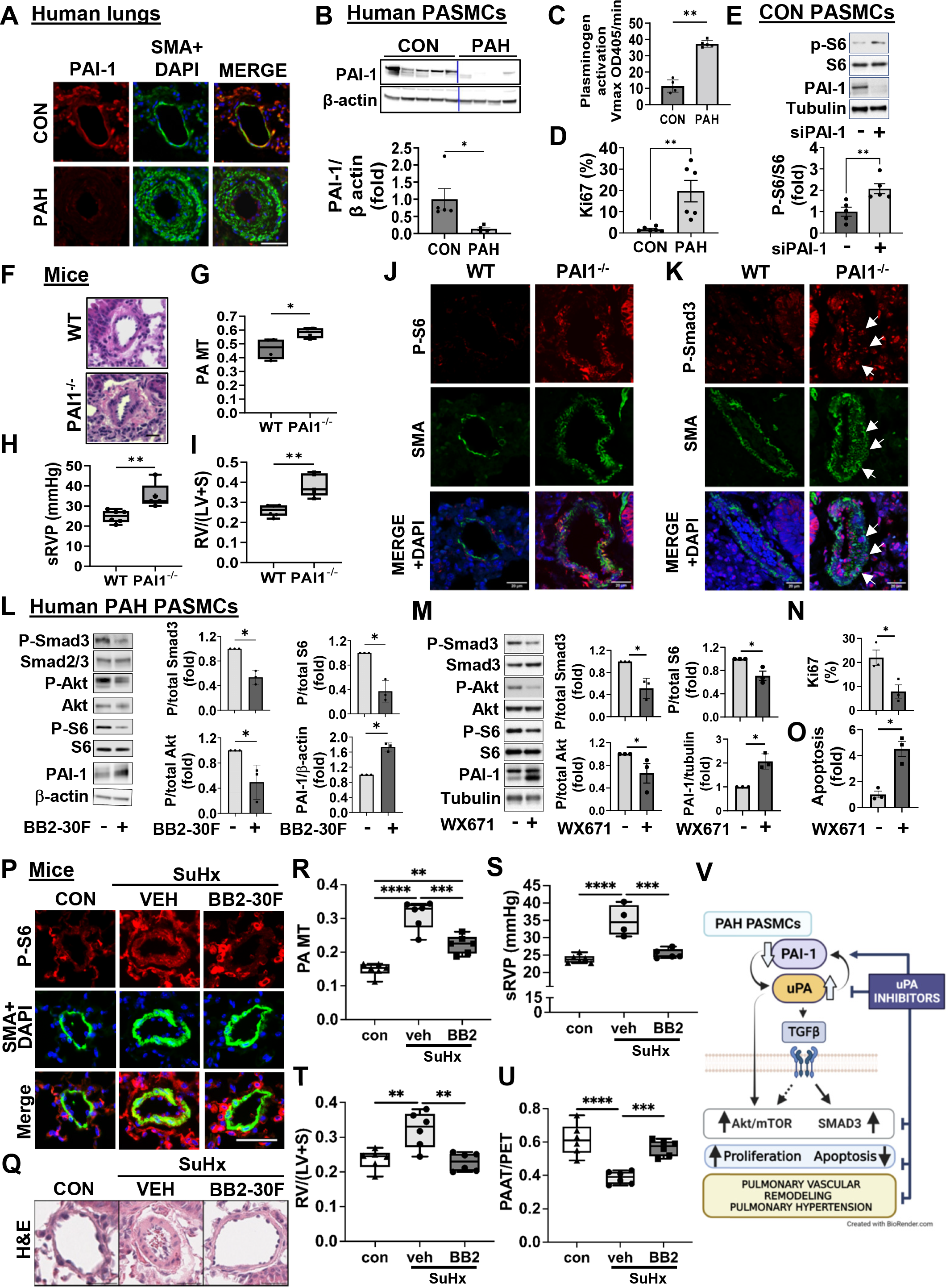
A: IHC analysis of human lung tissue sections to detect PAI-1 (red). Green - SM α-actin (SMA); blue - DAPI; yellow - red and green overlap. Images are representative from 3 subjects/group. Bar equals 50μm. **B:** Early-passage PASMCs from small (<1.5 mm outer diameter) muscular PAs from human control and PAH lungs were subjected to immunoblot analysis to detect indicated proteins. **C:** To measure plasminogen activation, 4 μM plasminogen and 1 mM chromogenic substrate for plasmin (S2251) were added to cells, and the kinetics of plasmin generation was measured at OD_405nm_. Rates of the plasmin generation were calculated and then normalized based on cell numbers/well. **D:** Cell proliferation was measured by staining of the cells for Ki67 and counting the percentile of Ki67-positive cells per total number of cells detected by DAPI. **B-D**: N=4-6 subjects/group. Data are means±SE. *p<0.05, **p<0.01 by Mann Whitney U. **E:** Human control PASMCs were transfected with siRNA PAI-1 (+) or control scrambled siRNA (-); 48h post-transfection, immunoblot analysis of whole cell lysates was performed to detect indicated proteins. Representative images (upper panel) and statistical analysis of n=3 subjects/group (lower panel) were shown. Data are means±SE, **p<0.01 by Mann Whitney U. **F-I:** Morphological and hemodynamic analysis of 10-month-old wild type C57BL/6J and PAI-1^-/-^ mice. **F,G**: Representative H&E images (bar equals 30μm) (**F**), PA medial thickness (PA MT) (**G**) from n=4 mice/group (2♂+2♀), 10 PAs (<150μm outer diameter) per mouse; **H, I:** sRVP (**H**) and RV/(LV+S) (Fulton index) (**I**) were measured in n=6 (3♂+3♀) control and n=5 (3♂+2♀) PAI-1^-/-^ mice. Data are means±SE; *p<0.05, **p<0.01 by Mann Whitney U. **J, K:** IF IHC analysis of lung tissue sections from PAI-1^-/-^ and WT mice. Red: phospho-Smad3 (**J**) or phospho-S6 (**K**), green - SMA, blue - DAPI. Bar equals 20 μm. Images are representative from 3 animals/group. **L, M:** Human PAH PASMC were incubated with 10 μM BB2-30F **(L)**, 10 μM WX671 **(M)** or diluent for 48 hr followed by immunoblot analysis to detect indicated proteins (**L, M**), cell proliferation (Ki67) (**N**), or apoptosis (*In Situ* Cell Death Detection Kit, Sigma) (**O**). Images are representative of three experiments, each performed using cells from different human subject. Data are means±SE, n=3 subjects/group; *p<0.05 by Mann Whitney U. **P-U:** Inhibition of uPA prevents development of PH in the SuHx mouse model. WT (C57BL/6J) male mice were given daily intraperitoneal injections of 10 mg/kg BB2-30F or vehicle from day 1-21 of exposure to SU5416 (20 mg/kg s.c. once a week) and chronic hypoxia (FiO_2_ 10%). Hemodynamic measurements and tissue harvest were made on day 21. Controls (CON) were same-age mice kept under normoxia. **P:** IF IHC of lung tissue sections. Red - phospho-S6, green - SMA, blue - DAPI. Bar equals 50 μm. Images are representative from 3 animals/group. **Q, R**: Representative H&E images (bar equals 30 μm (**Q**) and PA MT (**R**); n=6 mice/group, 10 PAs with outer diameter <150 μm per mouse. **S-U**: sRVP **(S)**, RV/(LV+S) (Fulton index) **(T)**, and PAAT/PET ratio **(U);** n=4-6 mice/group, **p<0.01, ***p<0.001,****p<0.0001. **V:** Schematic representation of the mechanism by which PAI-1 deficiency promotes pulmonary vascular remodeling and PH.

We then asked whether insufficiency of PAI-1 contributes to pulmonary vascular remodeling and PH *in vivo*. Histological examination of 10 month-old PAI-1 knock-out (PAI-1^-/-^) mice revealed remodeling of the small pulmonary PAs, characterized by an increase in medial wall thickness (PA MT) (**Fig. 1F, G**) accompanied by a significant increase in systolic right ventricular pressure (sRVP) (**Fig. 1H**) and right ventricle (RV) hypertrophy (Fulton index, RV/(left ventricle+septum, LV+S)) in PAI-1^-/-^ mice compared to age- and sex-matched wild type (WT) mice on the same background (**Fig. 1I**). These results demonstrate that loss of PAI-1 results in the development of spontaneous pulmonary vascular remodeling and PH.

We next investigated the mechanistic basis of pulmonary vascular remodeling and PH resulting from PAI-1 deficiency. uPA, unchecked in the absence of PAI-1, regulates vascular remodeling (reviewed in(8)) through plasmin-mediated activation of matrix metalloproteinases(9) and growth factors, including TGF-β1(9), which is over-produced by human PAH PASMC and promotes PASMCs hyper-proliferation, pulmonary vascular remodeling and PH via canonical Smad2/3 and non-canonical PI3K-Akt-mTOR pathways(10, 11). TGF-β1 is secreted by vascular cells and platelets in a biologically inactive (latent) form that must be proteolytically activated by uPA-generated plasmin to bind to their surface receptors(12). Based on previous work showing that mice deficient in uPA (*plau*^*-/-*^ mice) are protected from hypoxia-induced PH(13) and our observations that PAH PASMCs have a significant increase in plasminogen activation (**Fig. 1C**), we hypothesized that unchecked uPA activity promotes PA remodeling and spontaneous PH in PAI-1 knockout mice by activating TGF-β1 and mTOR pathways. Consistent with this hypothesis, immunostaining of lung sections of PAI-1^-/-^ mice showed increased phospho-Smad3 and phospho-S6 in small PAs compared to WT controls (**Fig. 1J, K**), suggesting that PAI-1 loss results in TGF-β/Smad3 and mTORC1 activation.

PAI-1 levels vary widely in response to diverse external stimuli. The synthesis and function of PAI-1 in the pulmonary vasculature and elsewhere are subject to complex regulatory pathways not currently amenable to control. Therefore, we asked whether the functional consequences of PAI-1 deficiency in pulmonary vasculature could be mitigated through control of uPA. We first examined the effect of amiloride analog BB2-30F, which inhibits both human and mouse uPA(14) and suppresses metastasis in mouse model of pancreatic cancer(15, 16), and WX671 (upamostat), an inhibitor of enzymatic activity of human uPA, which is currently in clinical trials for patients with pancreatic cancer(17), on human PAH PASMCs.

Addition of BB2-30F and WX671 to PAH PASMCs reduced TGF-β-dependent Smad3 phosphorylation and inhibited Akt-mTORC1 signaling, as evidenced by significant decreases in P-Smad3, P-S473 Akt, and P-S6 **(Fig. 1L, M)**. WX671 inhibited proliferation and induced apoptosis in human PAH PASMC (**Fig. 1N, O**). Of interest, inhibition of uPA activity was accompanied by an increase in the level of PAI-1 protein **(Fig. 1L, M)**. This suggests that excessive enzymatically active uPA further depletes PAI-1 in a feedforward pathogenic cycle and that inhibition of uPA may spare residual PAI-1 from further depletion in PAH PASMC.

Next, to test whether uPA mediate pulmonary vascular remodeling and PH *in vivo*, we examined the effect of inhibition of uPA catalytic activity by BB2-30F on the development of SU5416/Hypoxia (SuHx)-induced PH in mice. WT (C57BL/6J) male mice were given daily intraperitoneal injections of BB2-30F (10mg/kg) or vehicle from day 1-21 during exposure to SU5416 (20 mg/kg s.c. once a week) and chronic hypoxia (FiO2 10%). Hemodynamic measurements and tissue harvest were performed on day 21. Vehicle-treated SuHx-exposed mice had up-regulated mTORC1 (assessed by phospho-S6) in small PAs (**Fig. 1P**), developed pulmonary vascular remodeling and PH, as evidenced by significant increase of PA MT (**Fig. 1Q, R**), sRVP (**Fig. 1S**), RV hypertrophy (**Fig. 1T**), and a significant decrease in the pulmonary artery acceleration time/pulmonary ejection time (PAAT/PET) ratio (**Fig. 1U**) compared to age- and sex-matched normoxia controls, whereas BB2-30F-treated group was protected from all these pathological changes (**Fig. 1P-U**).

In conclusion, this study identifies an important role for PAI-1 deficiency and unrestricted uPA activity in PASMC remodeling and PH *in vitro* and *in vivo*, provides novel mechanistic link from PAI-1 through uPA-induced Akt/mTOR and TGFβ-Smad3 up-regulation to PASMC hyper-proliferation, remodeling, and PH (**Fig. 1V**), and suggests that inhibition of uPA to re-balance the uPA-PAI-1 tandem might provide a novel approach to complement current therapies used to mitigate this severe and progressive pulmonary vascular disease. These findings call for further investigation to determine potential attractiveness of this molecular pathway as a target system to reverse established PAH.

## ACKNOWLEDGEMENTS

This work is supported by NIH/NHLBI R01HL166932 (TVK), R01HL13026 (EAG), R01HL150638 (EAG), RO1HL141462 (VS), R01HL139881 (LF), RO1HL159256 (DBC), Nina Ireland Program for Lung Health (PJW), R35HL150698 (MP), NHMRC Ideas grant APP1181179 (MR and MJK). Pulmonary Hypertension Breakthrough Initiative is supported by NIH/NHLBI R24 HL123767 and by the Cardiovascular Medical Research and Education Fund (CMREF).

## REFERENCES

1. Johnson S, Sommer N, Cox-Flaherty K, Weissmann N, Ventetuolo CE, Maron BA. Pulmonary Hypertension: A Contemporary Review. Am J Respir Crit Care Med 2023.

2. Pullamsetti SS, Savai R, Seeger W, Goncharova EA. Translational Advances in the Field of Pulmonary Hypertension. From Cancer Biology to New Pulmonary Arterial Hypertension Therapeutics. Targeting Cell Growth and Proliferation Signaling Hubs. Am J Respir Crit Care Med 2017; 195: 425–437.

3. Wind T, Hansen M, Jensen JK, Andreasen PA. The molecular basis for anti-proteolytic and non-proteolytic functions of plasminogen activator inhibitor type-1: roles of the reactive centre loop, the shutter region, the flexible joint region and the small serpin fragment. Biol Chem 2002; 383: 21–36.

4. Conese M, Blasi F. Urokinase/urokinase receptor system: internalization/degradation of urokinase-serpin complexes: mechanism and regulation. Biol Chem Hoppe Seyler 1995; 376: 143–155.

5. Gupta KK, Donahue DL, Sandoval-Cooper MJ, Castellino FJ, Ploplis VA. Plasminogen Activator Inhibitor-1 Protects Mice Against Cardiac Fibrosis by Inhibiting Urokinase-type Plasminogen Activator-mediated Plasminogen Activation. Sci Rep 2017; 7: 365.

6. Chen T, Huang JB, Dai J, Zhou Q, Raj JU, Zhou G. PAI-1 is a novel component of the miR-17∼92 signaling that regulates pulmonary artery smooth muscle cell phenotypes. Am J Physiol Lung Cell Mol Physiol 2018; 315: L149–l161.

7. Kouri FM, Queisser MA, Königshoff M, Chrobak I, Preissner KT, Seeger W, Eickelberg O. Plasminogen activator inhibitor type 1 inhibits smooth muscle cell proliferation in pulmonary arterial hypertension. Int J Biochem Cell Biol 2008; 40: 1872–1882.

8. Tkachuk V, Stepanova V, Little PJ, Bobik A. Regulation and role of urokinase plasminogen activator in vascular remodelling. Clin Exp Pharmacol Physiol 1996; 23: 759–765.

9. Lijnen HR. Plasmin and matrix metalloproteinases in vascular remodeling. Thromb Haemost 2001; 86: 324–333.

10. Guignabert C, Humbert M. Targeting transforming growth factor-β receptors in pulmonary hypertension. Eur Respir J 2021; 57.

11. Kudryashova TV, Shen Y, Pena A, Cronin E, Okorie E, Goncharov DA, Goncharova EA. Inhibitory Antibodies against Activin A and TGF-beta Reduce Self-Supported, but Not Soluble Factors-Induced Growth of Human Pulmonary Arterial Vascular Smooth Muscle Cells in Pulmonary Arterial Hypertension. Int J Mol Sci 2018; 19.

12. Lyons RM, Gentry LE, Purchio AF, Moses HL. Mechanism of activation of latent recombinant transforming growth factor b1 by plasmin. Journal of Cell Biology 1990; 110: 1361–1367.

13. Levi M, Moons L, Bouche A, Shapiro SD, Collen D, Carmeliet P. Deficiency of urokinase-type plasminogen activator-mediated plasmin generation impairs vascular remodeling during hypoxia-induced pulmonary hypertension in mice. Circulation 2001; 103: 2014–2020.

14. N Ses, Buckley BJ, Jiang L, Huang M, Ranson M, Kelso MJ, Yu H. Disruption of Water Networks is the Cause of Human/Mouse Species Selectivity in Urokinase Plasminogen Activator (uPA) Inhibitors Derived from Hexamethylene Amiloride (HMA). J Med Chem 2022; 65: 1933–1945.

15. Buckley BJ, Majed H, Aboelela A, Minaei E, Jiang L, Fildes K, Cheung CY, Johnson D, Bachovchin D, Cook GM, Huang M, Ranson M, Kelso MJ. 6-Substituted amiloride derivatives as inhibitors of the urokinase-type plasminogen activator for use in metastatic disease. Bioorg Med Chem Lett 2019; 29: 126753.

16. Buckley BJ, Aboelela A, Minaei E, Jiang LX, Xu Z, Ali U, Fildes K, Cheung CY, Cook SM, Johnson DC, Bachovchin DA, Cook GM, Apte M, Huang M, Ranson M, Kelso MJ. 6-Substituted Hexamethylene Amiloride (HMA) Derivatives as Potent and Selective Inhibitors of the Human Urokinase Plasminogen Activator for Use in Cancer. J Med Chem 2018; 61: 8299–8320.

17. Heinemann V, Ebert MP, Laubender RP, Bevan P, Mala C, Boeck S. Phase II randomised proof-of-concept study of the urokinase inhibitor upamostat (WX-671) in combination with gemcitabine compared with gemcitabine alone in patients with non-resectable, locally advanced pancreatic cancer. Br J Cancer 2013; 108: 766–770.

